# Machine learning-based Personalized Dietary Recommendations to Achieve Desired Gut Microbial Compositions

**DOI:** 10.64898/2026.05.12.724618

**Authors:** Xu-Wen Wang, Dan Huang, Pingfeng Yu, Scott T. Weiss, Yang-Yu Liu

## Abstract

Dietary intervention is an effective way to alter the gut microbiome to promote human health. Yet, due to our limited knowledge of diet-microbe interactions and the highly personalized gut microbial compositions, an efficient method to prescribe personalized dietary recommendations to achieve desired gut microbial compositions is still lacking. Here, we propose a machine learning framework to resolve this challenge. Our key idea is to implicitly learn the diet-microbe interactions by training a machine learning model using paired gut microbiome and dietary intake data from a population-level cohort. The well-trained machine learning model enables us to predict the microbial composition of any given species collection and dietary intake. Next, we prescribe personalized dietary recommendations by solving an optimization problem to achieve the desired microbial compositions. We systematically validated this Machine learning-based Personalized Dietary Recommendation (MPDR) framework using synthetic data generated from an established microbial consumer-resource model. We then validated MPDR using real data collected from a diet-microbiome association study. The presented MPDR framework demonstrates the potential of machine learning for personalized nutrition.

## Introduction

The human gut microbiome plays a vital role in health, and its dysbiosis has been associated with many diseases^1–5^. Although many factors can alter the human gut microbiome, diet is known to be a major host factor shaping the human gut microbiome^6^. For instance, short-term high consumption of animal products can increase the abundance of bile-tolerant bacteria such as *Alistipes, Bilophila*, and *Bacteroides*, while decreasing the levels of fiber-degrading members of *Bacillota* (formerly *Firmicutes)*, such as *Roseburia, Eubacterium rectale*, and *Ruminococcus bromii*^7^, respectively. Long-term diets rich in animal fat, protein, processed foods, and sugar are associated with elevated levels of taxa (e.g., *Bacillota* and *Ruminococcus gnavus)* linked to pro-inflammatory activity, along with reduced abundances of Short-chain fatty acid (SCFA)-producing taxa such as *Bifidobacterium*^8^. This diet-microbe link implies the potential of using dietary intervention to effectively alter the gut microbiome to benefit host health. However, due to complex microbial interactions and highly personalized microbial compositions affected by many host factors (e.g., medication use, age, BMI, and geography^9–11^), the prescription of personalized dietary recommendations to achieve desired gut microbial compositions is still a very challenging task^12–15^.

Previous studies have used traditional machine learning algorithms to prescribe personalized dietary recommendations to achieve certain metabolic responses^16–18^. For example, leveraging blood parameters, dietary habits, and gut microbiome data, one can predict the postprandial glycemic response using a gradient-boosting regressor and design a personalized postprandial-targeting diet^16^. It has been shown that such a personalized postprandial-targeting diet improved glycemic control significantly more than the Mediterranean diet in pre-diabetes^18^. A recent pilot study used XGBoost (Extreme Gradient Boosting)for personalized dietary recommendations to improve symptoms of Irritable Bowel Syndrome (IBS), where the modulation of the gut microbiome leading to a decrease in the IBS severity index was identified^19^. Subsequently, the micronutrient composition that supports this modulation was calculated using the micronutrient database. We notice that personalized dietary recommendations in previous studies were all based on a single metabolite or a disease severity index, and the impact of the diet intervention on the gut microbiome itself was not explicitly considered.

Here, we propose a Machine learning-based Personalized Dietary Recommendation (MPDR) framework to achieve desired gut microbial compositions. This MPDR framework consists of two steps. First, we use paired gut microbiome and dietary intake data to train a machine learning model to implicitly learn the diet-microbe interactions. The well-trained machine learning model enables us to predict the gut microbial composition for any given microbial species collection and dietary intake. Second, we prescribe the personalized dietary recommendation by solving an optimization problem to identify the best dietary intervention to steer the gut microbiome of a given individual toward a desired gut microbial composition. We systematically validated this MPDR framework using both synthetic and real data.

## Results

### The MPDR framework

We consider the human gut microbiome as a meta-community that harbors a pool of *N* different microbial species. Denote the species pool of this meta-community as Ω = {1, ⋯, *N*}, and a set of gut microbiome samples as 𝒮 = {1, …, *S*} collected from *S* individuals. The gut microbiome of any individual can be viewed as a local community assembled from a particular subset of species in Ω. Before any dietary intervention, the species assemblage of an individual’s baseline gut microbiome is represented by a binary vector ***z*** ∈ {0,1}^*N*^. For a given dietary intervention, the dietary intake is represented by a vector 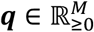 with *M* the total number of food items.

The MPDR framework consists of two steps. First, we implicitly learn the diet-microbe interactions by training a machine learning model (**Fig.1a)**. This is achieved by learning a map from the species assemblage ***z*** of an individual’s baseline gut microbiome and the dietary intake profile ***q*** to their gut microbial composition ***p*** ∈ Δ^*N*^, i.e., *φ*(***z, q***) ↦ ***p***. Here 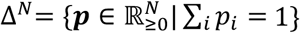. Note that we assume this map is unique. In other words, given a (***z, q***) pair, there is a unique ***p***. However, given a species assemblage ***z***, there could be multiple ***q*** vectors that will yield the same ***p*** vector. Here, we employ a classical machine learning model, Multi-Layer Perceptron (MLP), to learn the map *φ* (see **Methods)**, motivated by its superiority in computational efficacy and its flexibility in managing the dimensionality difference between the nutrient profile and microbial profile^20^. When provided with a baseline gut microbial assemblage ***z*** and dietary intake profile ***q***, this map *φ* will enable us to predict the corresponding gut microbial composition ***p***.

**Figure 1:**
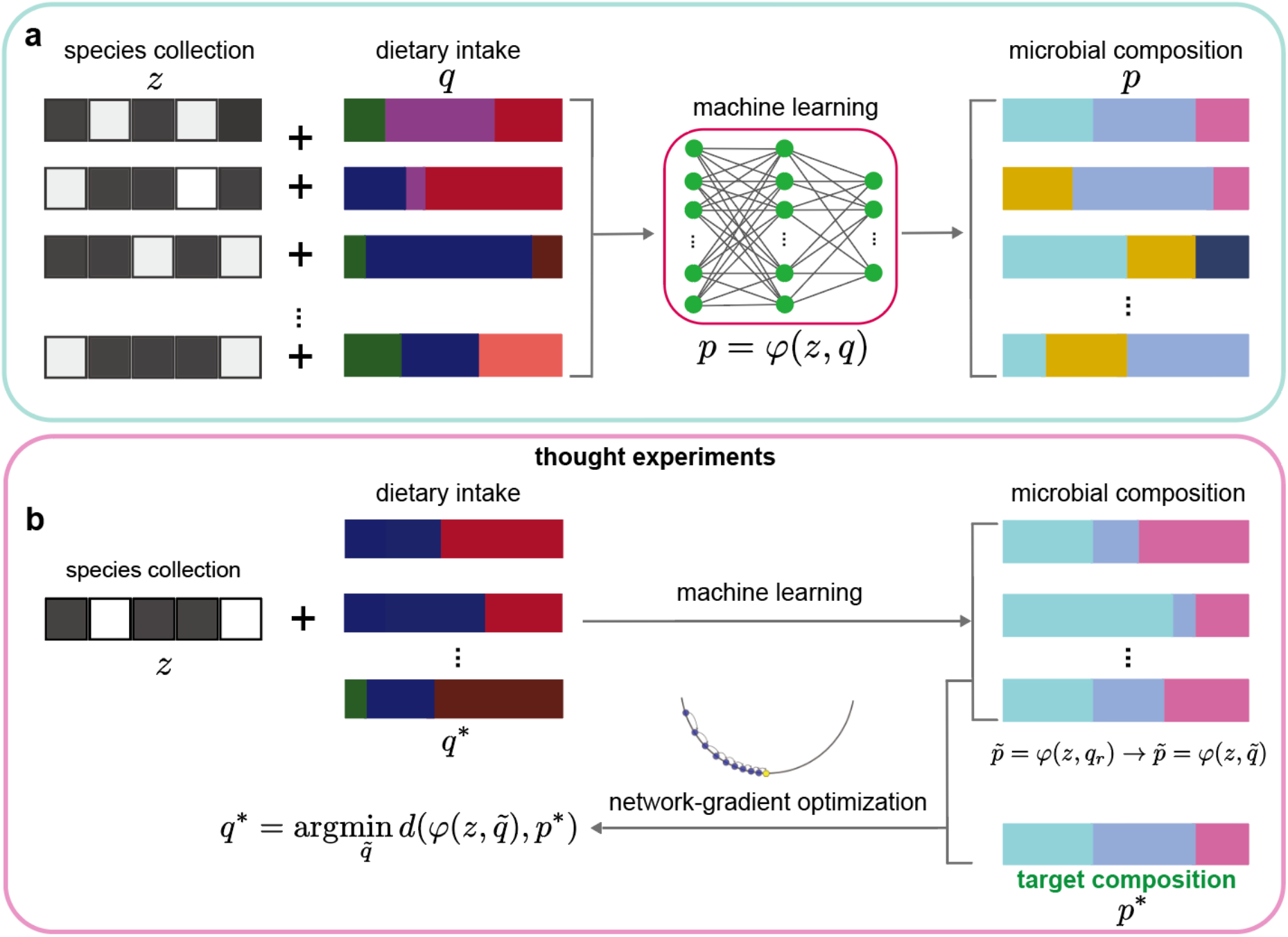
Workflow of the Machine learning-based Personalized Diet Recommender (MPDR) framework. **a**, The baseline species assemblage of a microbiome sample *s* is represented by a binary vector ***z*** ∈ {0,1}^***N***^, where its *i*-th entry satisfies *z*_*i*_ = 1 (*z*_*i*_ = 0) if species-*i* is present (or absent) in this sample. The dietary intake of this sample is characterized by a vector ***q***, where its *i*-th entry ***q***_*i*_ is the abundance of nutrient-*i*. The microbial composition of this sample is characterized by a vector ***p*** ∈ Δ^***N***^, where its *i*-th entry ***p***_*i*_ is the relative abundance of species-*i* in this sample and Δ^***N***^ is the probability simplex. The machine learning model is trained to learn the map (***z*** ∈ {0,1}^***N***^, ***q***) *↦* ***p*** ∈ Δ^***N***^. **b**, We conduct a *thought regulation* by revising the dietary profile ***q*** for any new sample to obtain a new dietary profile ***q***_P_. Then, for the new species collection and dietary profile (***z, q***_*r*_), we use machine learning model to predict its new composition 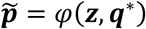. The optimal dietary profile is obtained by an optimization algorithm to minimize a loss function, e.g., the lowest dissimilarity with the compositions of a target population.

Second, to prescribe a personalized dietary recommendation to individual *s* (with a baseline or pre-intervention gut microbial species assemblage ***z)*** to achieve their desired post-intervention gut microbial composition ***p***\*, we conduct a series of “thought” dietary interventions to identify the optimal diet ***q***\* (**Fig.1b)**. Each thought dietary intervention is generated by updating a set of logits. For a thought dietary intervention that results in a new diet 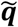, we use the well-trained MLP to predict the recommended-diet projected microbial composition 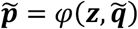. To find the optimal diet ***q***\*, we choose the Bray-Curtis dissimilarity 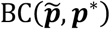 between the recommended-diet projected composition 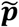 and the target microbial composition ***p***\* as the loss function *L* and try to minimize it. In other words,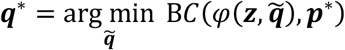. To solve this optimization problem, we apply gradient-based optimization to the logits parameterization of the diet (**Fig.1b**, and **Methods**).

Note that we can define the target microbial composition ***p***\* in many different ways. For instance, we can define it to be the baseline healthy microbial composition of an individual. We can also define it to be the composition that has the shortest average dissimilarity to the microbial composition of each individual in a healthy population, i.e., the geometric median of the healthy microbial compositions. We can also define the loss function *L* to be the negative relative abundance of a specific beneficial species that needs to flourish in the gut (e.g., those species that can produce short-chain fatty acids^21–25^), thereby allowing the optimal diet to boost its growth.

We emphasize that here, pre-intervention and post-intervention simply refer to the baseline and follow-up states rather than causal contrasts. The dietary recommendations generated by MPDR should be interpreted as computationally optimized inputs under learned associations, not as interventions with guaranteed causal effects. To establish causal effects of dietary changes on microbiome structure, experimental validation or causal modeling would be required.

### Validation of MLP in microbial composition prediction

The machine learning model MLP is a key component of the MPDR framework. To validate MLP in the prediction of microbial composition from paired species assemblage and dietary intake profile, we generated synthetic data using the Microbial Consumer Resource Model (MiCRM)^26^ with *N* = 100 species in the species pool and *M* = 60 resources (**Methods**). MiCRM describes the per-capita growth of species as a function of the resource consumption rate. Each species *i* is characterized by a resource utilization rate *U*_*iα*_, i.e., the rate species *i* consumes the resource *α*. The entries of the resource utilization matrix *U* = (*U*_*iα*_) were initialized using a uniform distribution with a parameter *g* ∈ {0.01, 0.1, 0.5} to control the specialists/generalists ratio. We also introduced a parameter *C* ∈ {0.2, 0.4, 0.6} to control the probability of a species *i* consuming a resource *α*.

To initialize each sample (i.e., each local community), we selected 80 species from the species pool (*N* = 100). Among the 80 species, 50 species were common across all samples, and the remaining 30 species were randomly drawn from the other 50 species in the pool. We initialized the resource profile by randomly generating resource abundances from a uniform distribution. The difference in the initial resource profiles across different samples was controlled by a resource variability parameter *σ* ∈ {0.02, 0.05, 0.1}. A lower *σ* value represents greater universal resource profiles. The *σ* is set to be about 0.05 (**Fig.S1**), so that the median dissimilarity among all resource profiles is comparable with that in real data^6,27^.

For each sample, starting from its initial species collection and resource profile, we first ran MiCRM to reach a pre-intervention steady state with species collection denoted by ***z***. Then, we simulated a dietary intervention to the steady state by perturbing its resource profile ***q*** and ran MiCRM again to reach another steady state, representing the post-intervention or final state ***p***. We generated *S* samples this way. The species collection of the initial steady state ***z***, the resource profile ***q***, and the microbial composition ***p*** of the final state of the *S* samples were used to train MLP to minimize the masked Dirichlet negative log-likelihood loss between the true and predicted microbial compositions (see **Methods**). To investigate how training sample size affects prediction accuracy, we systematically varied the ratio of sample size to species richness 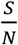 across the values 0.5, 1, 2, 5, and 10. We also generated additional samples in the same way as the test data to evaluate model performance.

To systematically evaluate the performance of MLP on test data generated using MiCRM with different ecological parameter settings (*σ, C, g*), we quantified prediction errors using Bray-Curtis dissimilarity and Aitchison distance between the true and predicted microbial compositions. The prediction error decreased consistently as the training sample size relative to the number of species (*S*/*N*) increased, reaching a plateau around (*S*/*N*)= 5 (**Fig.2a-c**). The minimal Bray-Curtis dissimilarity is lower than 0.1 for small variation *σ* = 0.02 (**Fig.2a**). In addition, we found that the prediction error is relatively large for higher *C* (**Fig.2b**), due to more complex resource-species interactions. Moreover, increasing the consumer specialism level *g* reduces the prediction error (**Fig. 2c**), because a larger *g* (*g* = 0.5) makes species’ resource preferences more distinct, simplifying the resource-species relationships that the model needs to learn. Note that this result is robust using Aitchison distance (see **Fig.S2**). Furthermore, Pearson correlation coefficients (*R*) between the predicted and true relative abundances demonstrated high predictive accuracy across all parameter sets (*σ* (**Fig.2d**), *C* (**Fig.2e**), and *g* (**Fig.2f**)), respectively. As *S*/*N* increased, *R* rapidly approached 1.0, confirming the strong agreement between model predictions and ground-truth abundances. Finally, direct comparisons of different representative training sizes (*S*/*N* = 1, 5, 10, respectively) revealed that MLP can accurately predict species’ relative abundances if the sample size is large enough (**Fig.2g-i**): *R* > 0.97 with *p* − value < 0.001 (two-sided t-test) for *S* = 5*N*. Moreover, we interrogated the MLP architecture itself by comparing single- and two-layer models with varying hidden units. We found that two-layer models consistently outperformed single-layer models. For two-layer models, increasing the number of hidden units beyond 256 provided minimal benefit (see **Fig.S3**).

**Figure 2:**
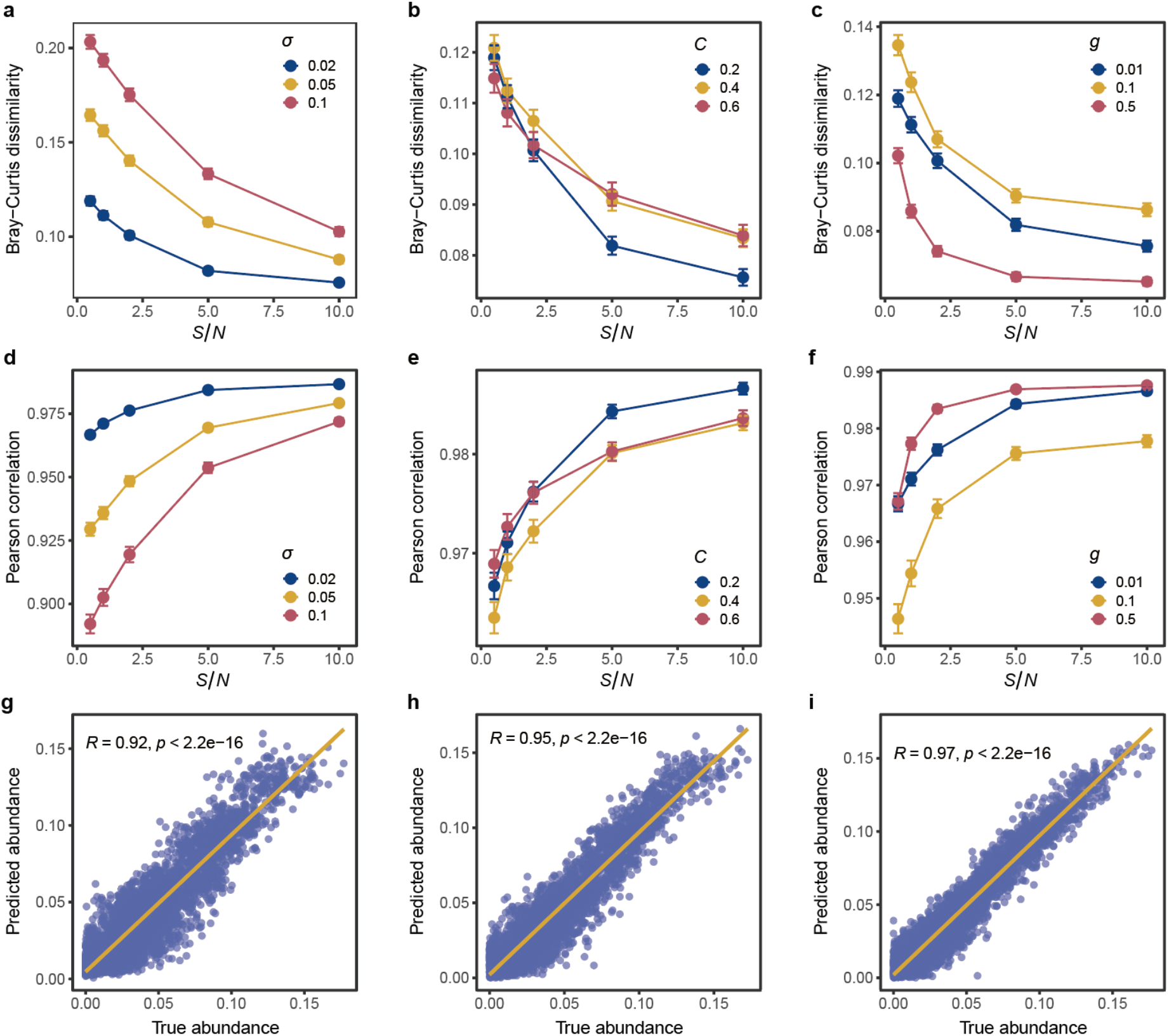
In silico validation of microbiome composition prediction in MPDR framework. Results are obtained for the pools of *N* = 100 species and 60 resources with a microbiome consumer-resource model. We generated *S* samples (50-1000) by a microbiome consumer-resource model with different nutrition variability *σ*, connectivity *C*, and specialist *g*, to train MPDR. **a-c**, Prediction error quantified as Bray-Curtis dissimilarity between the true compositions and predicted compositions. The error bar represents the mean of the standard deviation over 10 random splitting datasets. In panel **a**, *C* = 0.2 and *g* = 0.01. In panel **b**, *σ* = 0.02 and *g* = 0.01. In panel **c**, *σ* = 0.02 and *C* = 0.2. **d-f**, Prediction error quantified as the Pearson correlation coefficient *R* between the true and predicted relative abundances. In panel **d**, *C* = 0.2 and *g* = 0.01. In panel **e**, *σ* = 0.02 and *g* = 0.01. In panel **f**, *σ* = 0.02 and *C* = 0.2. **g-i**, Predicted versus true microbial compositions with increasing training sample size. Here, *S*denotes the baseline training size, *N* = 100 samples, and 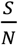 is the ratio between the number of training samples and the number of species. Thus, panels (**g-i**) correspond to 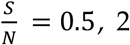, and 5 (i.e.,training sample sizes of 50, 200, and 500), respectively. Parameter settings: *σ* = 0.02, *C* = 0.2 and *g* = 0.01.

We further benchmarked the baseline MLP model against a Transformer-based architecture (see **Methods**) that retained an identical MLP prediction head, as well as three classical regression baselines, multi-output Support Vector Regression (SVR), Random Forest (RF) regression, and XGBoost regression (**Fig.S4**). The Transformer model encodes diet and baseline microbiome information using two independent Transformer encoders with tokens globally reordered by abundance, thereby imposing an explicit core-to-rare structure prior to feature fusion^28^. Across almost all the training sample sizes, the MLP consistently achieved the lowest Bray-Curtis dissimilarity (**Fig.S4a**,**e**,**i**,**m**) and RMSE (**Fig.S4c**,**g**,**k**,**o**), as well as the highest Pearson correlation coefficients (**Fig.S4b**,**f**,**j**,**n**) and R-square (**Fig.S4d**,**h**,**l**,**p**), indicating superior prediction accuracy and stability. Among the classical baselines, XGBoost achieved the best performance, RF obtained intermediate accuracy, and SVR showed the poorest predictive power (**Fig.S4)**. However, none of these methods matched the robust performance of the MLP, which remained the top-performing model across almost all settings. These findings suggest that, despite its simpler architecture, the MLP captures the nonlinear mapping between diet profiles and microbial compositions more effectively than the more complex transformer-based frameworks.

### Validation of MPDR in microbiome regulation

Next, we validated the performance of MPDR in diet-based microbiome regulations. To simulate the microbial compositions associated with different dietary patterns, we first generated 200 baseline microbiome samples using MiCRM as described in the previous subsection. The microbial composition of each sample represents a desired target microbial composition ***p***\*. Then, for each sample, we randomly perturbed its resource profile ***q***_H_ to get an “undesired” resource profile ***q***_U_ and re-ran MiCRM to get the perturbed and undesired microbial composition ***p***_U_ (see **Methods**). Given the undesired resource profile ***q***_U_, the initial species assemblage ***z***, and the target microbial composition ***p***\*, we used MPDR to prescribe the personalized optimal diet ***q***\* to drive the microbiome towards the target composition ***p***\* as closely as possible. To examine whether the recommendation prescribed by MPDR works, we ran MiCRM again by using ***q***\* as the resource profile to obtain the new or recommended-diet projected composition 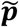. We found that indeed 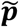 was much closer to ***p***\* than the undesired composition ***p***_U_ in the PCoA plot (**Fig.3a**). To further demonstrate this result, for a given sample, we compared the dissimilarity between its undesired composition ***p***_U_ (or its recommended-diet projected composition 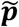) and its target composition ***p***\*, denoted as *d*(***p***_U_, ***p***\*) and 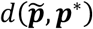, respectively (**Fig.3b**). We found that 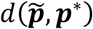 is much lower than *d*(***p***_U_, ***p***\*), meaning that 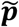 is much more similar to ***p***\* than ***p***_U_, regardless of connectivity *C* = 0.2 (**Fig.3**) or *C* = 0.4 (**Fig.S5**).

**Figure 3:**
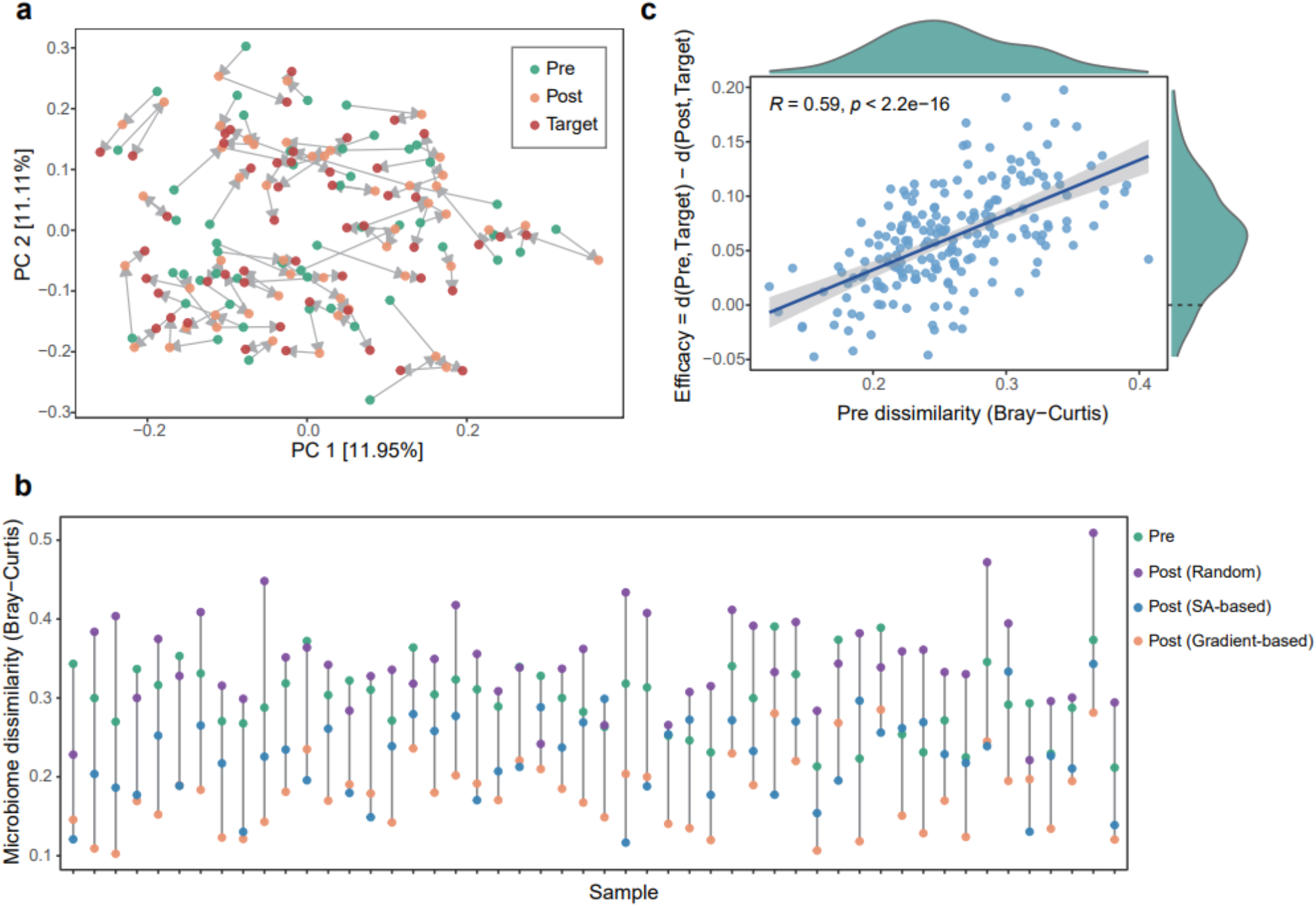
In silico validation of dietary recommendation in MPDR framework. Results are obtained for the pools of *N* = 100 species and 60 resources with a microbiome consumer-resource model. We generated 500 samples to train MPDR and an additional 200 samples to validate the recommendation of MPDR. For each validation sample with target microbial composition ***p***^*^, we randomly perturbed its nutrient profile ***q***_*H*_ into another nutrient profile ***q***_U_, and associated baseline microbial composition ***p***_U_,. MPDR was applied to find the optimal nutrient profile ***q***^*^ that minimizes dissimilarity between recommended-diet projected composition 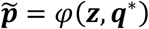 and ***p***^*^. Then, ran MiCRM again by using ***q***^*^ as the nutrient profile to obtain the new or recommended-diet projected composition 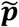, PCoA plot showing the microbial composition dissimilarity between baseline compositions (***p***_*U*_), recommended-diet projected 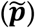 and target compositions (***p***^*^). **b**, Microbiome dissimilarity (Bray-Curtis) between the baseline microbiome composition (***p***_U_,) and the target composition (***p***^*^) (Pre, green), and between the post-intervention microbiome compositions 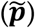 obtained using different diet recommendation strategies: random, simulated annealing (SA), and gradient-based optimization, and the target composition (***p***^*^)(Post). **c**, Recommendation efficacy 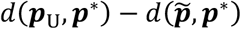 versus the pre-dissimilarity *d*(***p***_U_,, ***p***^*^). In panels a-c, *σ* = 0.1, *C* = 0.2 and *g* = 0.01. In panels a and b, we only showed 50 samples with the highest recommendation efficacy.

We defined the efficacy of the diet recommendation as 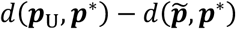, finding that the efficacy was strongly positively correlated with *d*(***p***_U_, ***p***\*) (**Fig.3c, Fig.S5)**. In other words, the higher the dissimilarity between the undesired microbiome ***p***_U_ and the target microbiome ***p***\*, the higher the efficacy of the dietary recommendation prescribed by MPDR. This result also implied that if the undesired composition is already extremely similar to the target composition, it becomes very challenging to further alter the microbiome through dietary intervention. Note that, we can always prescribe an optimal diet ***q***\*to reach the target microbial composition ***p***\*. However, due to the presence of prediction error in learning mapping *φ*, the true recommended-diet projected composition could still be slightly different from the desired composition.

Instead of community-wised regulation, we also validated MPDR in boosting the relative abundance of a single species. We defined the objective function as *L* ≡ −*x*_*t*_, where *x*_*t*_ is the relative abundance of the target species. We chose the species with the highest prevalence in the population as target species. We found that the relative abundance of the target species in the recommended-diet projected sample is higher than its original relative abundance in baseline community (**Fig.4a**). Moreover, the recommendation efficacy (defined as the difference between the relative abundances of target species in recommended-diet projected and baseline community) is significantly correlated with the pre-abundance of the target species (**Fig.4b**). This means that, if the relative abundance of target species is already extremely high, it becomes very challenging to further increase its abundance through dietary intervention. These results held, regardless of connectivity *C* = 0.2 (**Fig.4a,b**) or *C* = 0.4 (**Fig.4c,d**).

Moreover, we compared our gradient-based (GB) diet recommendation with simulated annealing (SA, see **Methods)** and random recommendation strategies. The results show that the GB method consistently achieves superior performance at both the community (**Fig.3b** and **Fig.S5b**) and single-species levels (**Fig.4a,c**), followed by SA, with random recommendations performing worst. This comparison demonstrates that MPDR provides effective and non-trivial diet recommendations.

**Figure 4:**
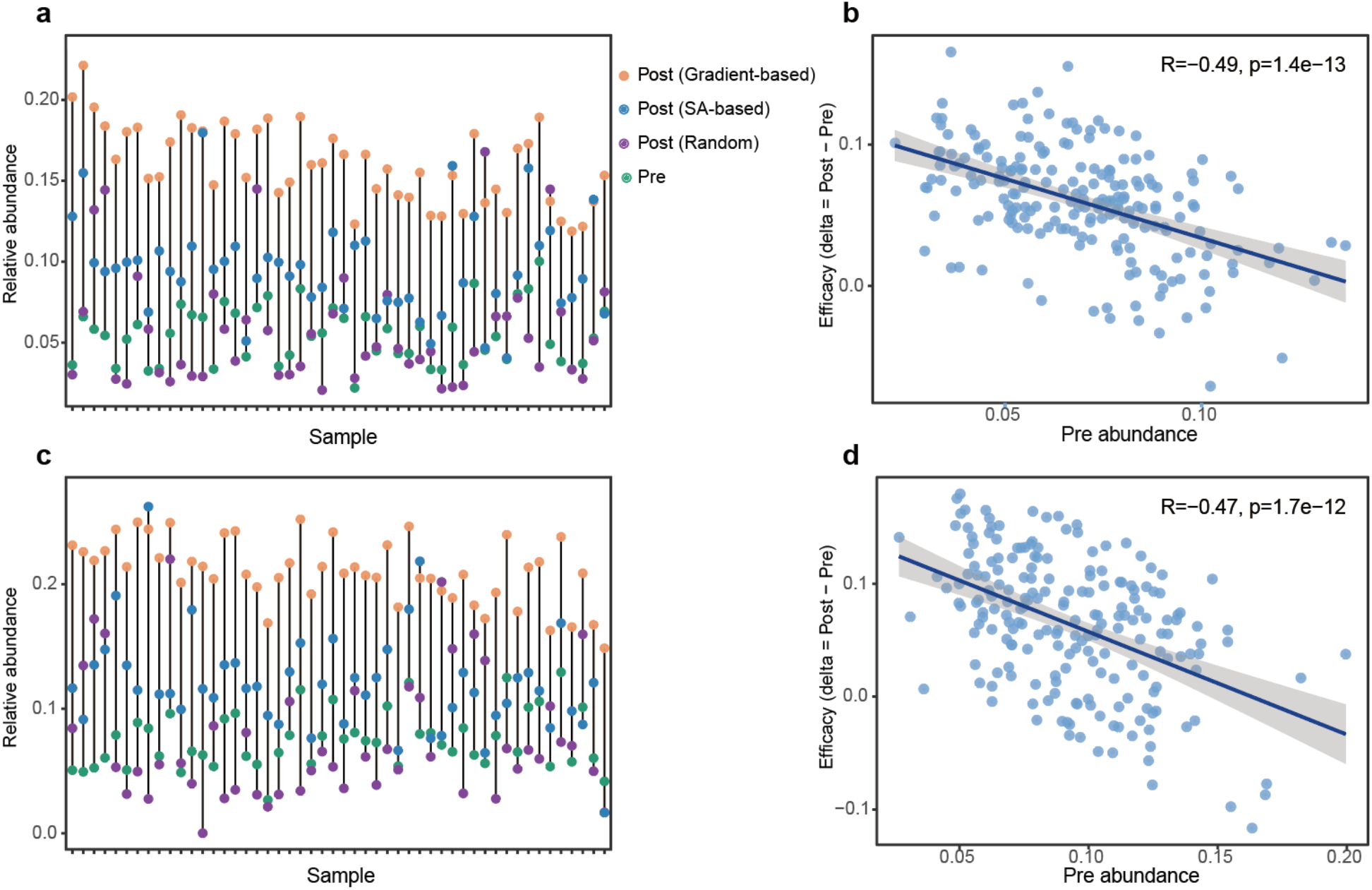
In silico validation of dietary recommendation in MPDR framework for species-wised regulation. Results are obtained for the pools of *N* = 100 species and 60 resources with a microbiome consumer-resource model. We generated 500 samples to train MPDR and an additional 200 samples to validate the recommendation of MPDR. For each validation sample, we randomly perturbed its nutrient profile ***q***_*H*_ into another nutrient profile ***q***_*U*_, and associated baseline microbial composition ***p***_*U*_,. MPDR was applied to find the optimal nutrient profile ***q***^*^ that minimizes the relative abundance of target species *x*_*t*_. Then, ran MiCRM again by using ***q***^*^ as the nutrient profile to obtain the new or recommended-diet projected composition 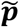 Relative abundance of target species in baseline composition ***p***_U_, (pre, green) and recommended-diet projected composition 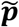 obtained using different diet recommendation strategies: random, simulated annealing (SA), and gradient-based optimization, and the target composition (***p***^*^)(Post). **b**,**d**, Recommendation efficacy 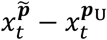 versus the pre-relative abundance of target species 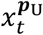.In panels **a** and **b**, *σ* = 0.1, *C* = 0.2 and *g* = 0.01. While in panels **c** and **d**, recommendations by MPDR with *C* = 0.4. In panels **a** and **c**, we only showed 50 samples with the highest recommendation efficacy.

### Application of MPDR in human gut microbiome regulation

Next, we applied the MPDR framework to the Diet-Microbiome Association Study (DMAS)^6^: a longitudinal study of 34 healthy individuals with dietary intake data and stool samples collected daily over 17 consecutive days. Daily dietary intake data was collected using the automated self-administered 24-h (ASA24) dietary assessment tool^29–31^. DMAS represented diet directly at the food level using the Food and Nutrient Database for Dietary Studies (FNDDS) food coding scheme^32^. The dietary input comprised 9 food groups (e.g., grains, fruits, and vegetables) encompassing 1,245 individual food items, together characterized by 65 nutrient attributes derived from their FNDDS nutrient profiles^32^. Both model training and dietary recommendation were conducted in the food space (i.e.,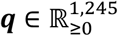). Microbiome features were filtered based on prevalence: species present in more than 15_*i*_ of samples were retained, resulting in 138 species and 50 genera. As the DMAS dataset did not directly involve any dietary recommendation or intervention, we analyzed this data using cross-validation manner. We randomly selected 283 samples from 25 individuals to train the MLP model, and we used the remaining 107 samples from the remaining 9 individuals to test it. We considered the species assemblage ***z*** at day *t* as baseline composition and microbial composition at day (*t* + 1) as the target compositions ***p***\*. For each test sample, we generated a randomized null (or undesired) composition ***p***_U_ by retraining the same species assemblage ***z*** while drawing species abundances from a uniform distribution over [0,1], followed by normalization to sum to 1. Then, we generated the randomly perturbed diet ***q***_U_ by randomly shuffling each individual’s consumption among consumed food items in DMAS. We then applied MPDR to make recommendations to get the optimal diet ***q***\*, aiming to steer the perturbed/undesired composition ***p***_U_ to the target compositions ***p***\*. To ensure each sample’s dietary pattern remains largely unchanged, we restricted the total grams of the recommended diet to match the pre-intervention diet.

Principal Coordinate Analysis shows that there is a large difference between baseline (green)and target compositions (red) (see **Fig.5a**). Then, we applied MPDR to prescribe diet recommendations, finding that the MLP-predicted post-intervention compositions (yellow) became much closer to the target compositions (red) (see **Fig.5a**). This clearly demonstrated that MPDR can indeed prescribe personalized dietary recommendations to approach desired gut microbial compositions. We found that, in contrast to randomly perturbed dietary intake without adherence to any dietary patterns (**Fig.5b**), the diet recommended by MPDR (**Fig.5c**) closely aligns with the real diet patterns (**Fig.5d**). For instance, the most prevalent and abundant food items are water, tea and coffee. Furthermore, for any given sample, we found that the food items prescribed by MPDR encompass multiple food groups, including beverages, fruits, dairy, and vegetables, thereby reflecting a more structured and diverse dietary composition. We also computed the diet dissimilarity between each test sample’s recommended (random)-food profile and its real food profile (**Fig.5e**). We found that recommended-food profile is closer to the real food profile than random-food profile (p<2.2e-16, Wilcoxon test).

**Figure 5:**
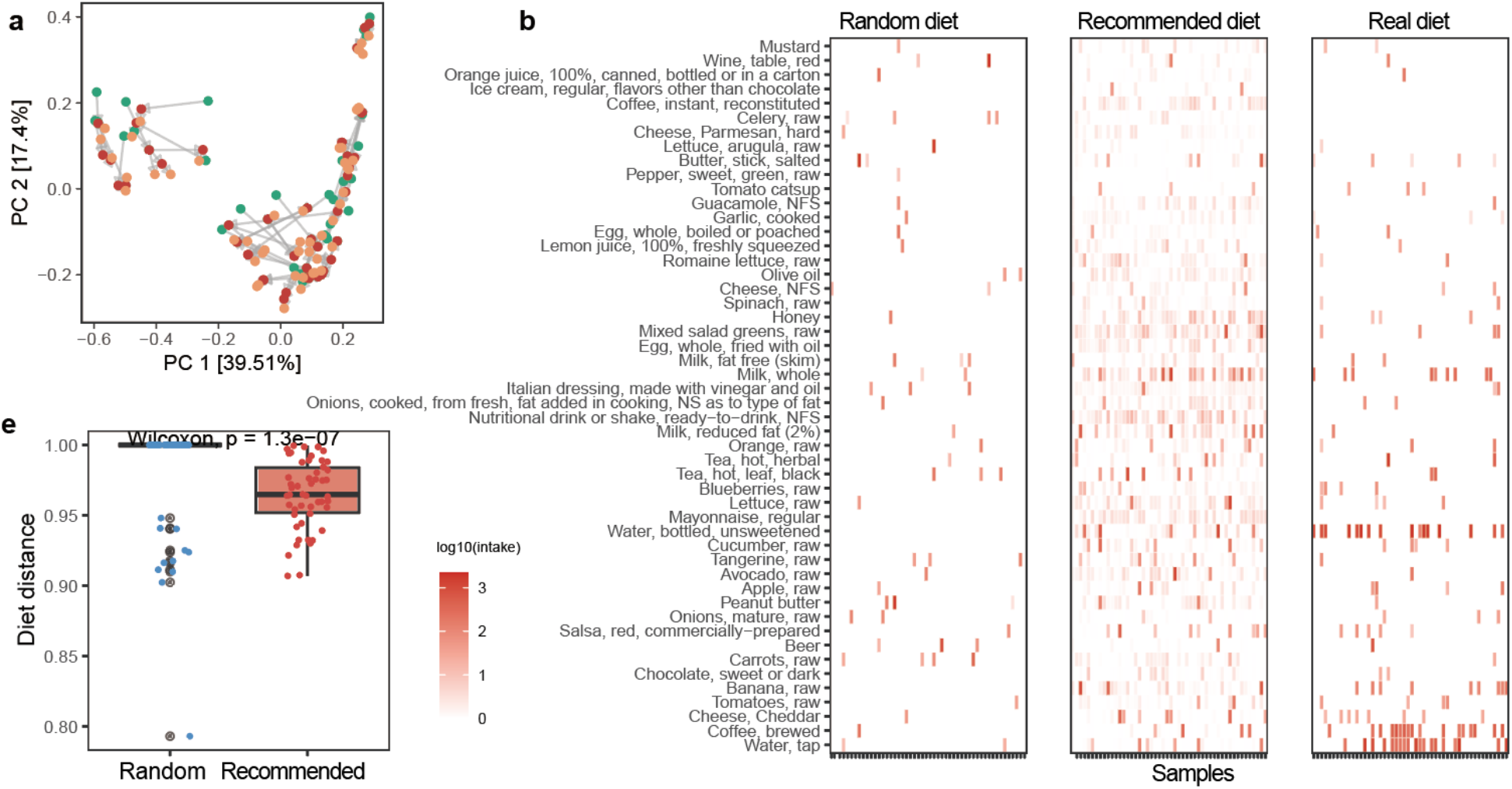
*In vivo* validation of the MPDR framework for community-wised regulation at species-level. We applied the MPDR framework into a Diet-microbiome association study (DMAS)^6^: a longitudinal study of 34 healthy individuals with dietary intake data and stool samples collected daily over 17 consecutive days. We randomly selected 25 individuals to train the MPDR framework, and the remaining 9 individuals were used to evaluate (in total 107 samples). We generated a random randomly perturbed composition ***p***, (pre) for each test sample with the same species assemblage ***z***, but the relative abundances were drawn from a uniform distribution. **a**, Principal coordinate analysis showing the microbiome communities of baseline (pre, green), recommended-diet projected (post, orange), and target compositions (target, red). For visualization purposes, we only showed top 50 samples with the highest regulation efficacy. **b-d**, Randomly perturbed food profiles (**b**), recommended (**c**) prescribed by MPDR and real food profiles (**d**). We only showed 50 most prevalent foods and top 50 samples with the highest regulation efficacy for visualization purposes. **e**, Diet dissimilarity between each test sample’s random or recommended-food profile and its real food profile. P-value was computed by paired Wilcoxon signed-rank test.

To identify those taxa that most strongly contributed to the composition predictions, we ranked taxa using a relative prediction error metric (mean absolute error normalized by each taxon’s mean baseline abundance), excluding extremely low-abundance taxa (< 0.001). As shown in **Fig.S6**, the taxa with the lowest relative prediction error were primarily members of *Alistipes, Bacteroides*, and *Roseburia*. These genera are well-known to be highly diet-responsive in human and animal studies, especially in relation to dietary fiber, protein, fat, and short-chain–fatty-acid metabolism^7,33,34^. The convergence between our model’s most accurately predicted taxa and those consistently identified as diet-sensitive supports the biological relevance of the diet-microbe relationships captured by our framework.

We further validated the performance of MPDR at the genus level. As shown in **Fig.S7a**, genus-level modeling achieved significantly lower Bray-Curtis dissimilarities than species-level modeling during the MLP training stage, reflecting reduced sparsity and increased functional coherence within genera. Importantly, this advantage also persisted in the dietary recommendation stage (**Fig.S7b**) where the Bray-Curtis dissimilarity between the desired microbial composition and the model-predicted endpoints under recommended diets was likewise reduced. Together, these results demonstrate that MPDR remains effective across different taxonomic resolutions. While species-level modeling enables fine-grained interpretation when well-characterized taxa are detectable, genus-level aggregation captures a broader fraction of the gut microbial community and more robustly reflects community-level functional responses, making genus-level MPDR a scalable and holistic framework for dietary regulation.

### Inferring diet-microbe interactions

To interpret MPDR, we proposed a susceptibility analysis to infer diet-microbe interactions. For each sample, we perturbed resource *α* by adding a small amount Δ*r*_*α*_ to its abundance *r*_*α*_ while holding all other resources fixed, and then used the well-trained MLP to re-predict the microbial composition to quantify each species’ response (i.e., its relative abundance change, denoted as Δ*p*_*i*_ for species *i*) to the perturbation of resource *α*. For each training sample, we repeated this procedure for each resource, yielding a sample-specific susceptibility matrix *S* = (*S*_*iα*_), with *S*_*iα*_ ≡ Δ*p*_*i*_/Δ*r*_*α*_ (see **Fig.6a**). The final susceptibility matrix was obtained by averaging the sample-specific *S* matrix across all samples, providing a population-level estimate of species-resource consumption interactions. Positive or negative *S*_*iα*_ indicates that increasing the abundance of resource *α* promotes or suppresses species *i*, respectively.

**Figure 6:**
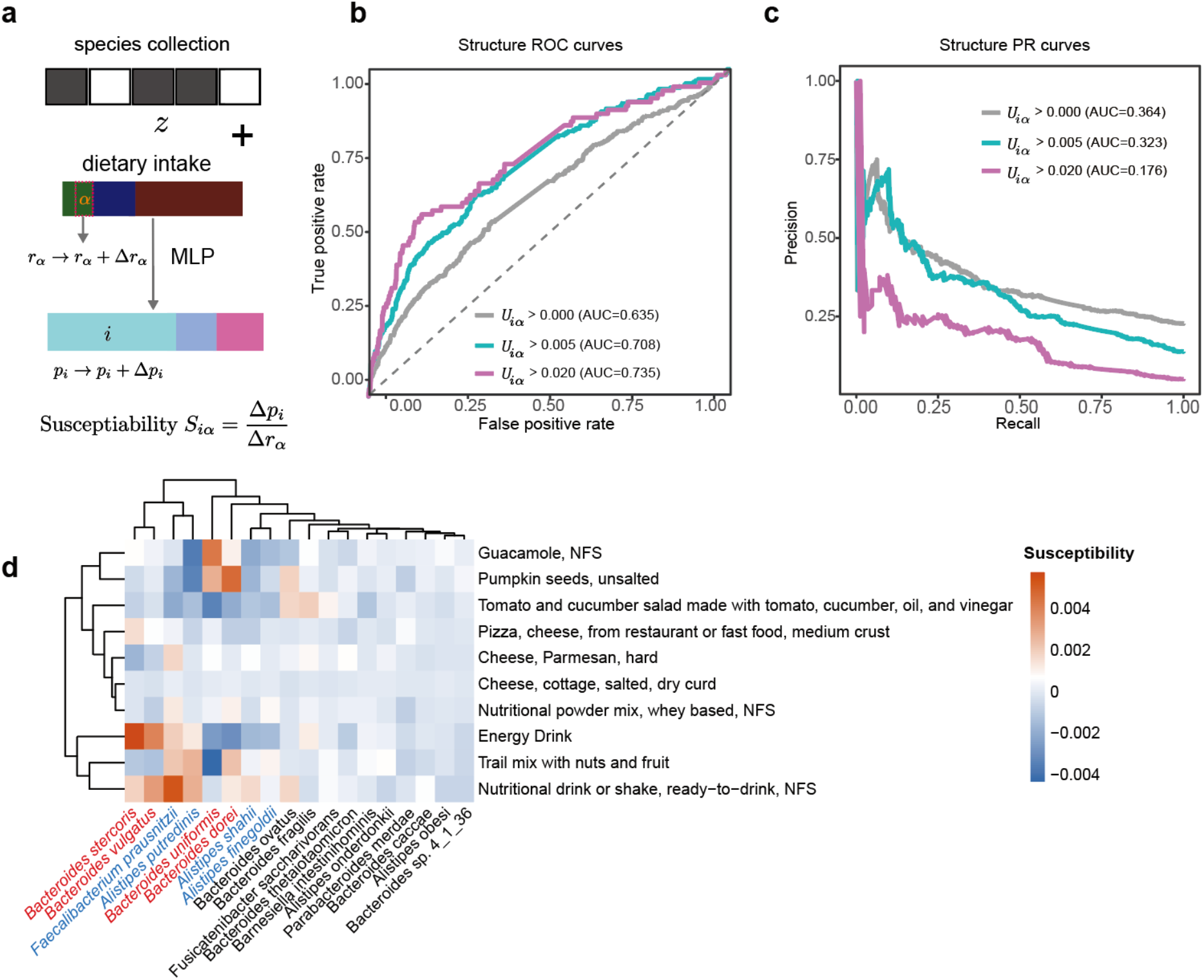
Using susceptibility of microbial compositions to diet perturbation of a well-trained MLP to infer diet-microbe interactions in both synthetic and real data. **a**, The susceptibility of the relative abundance (*p*_*i*_) of species *i* to the amount (*r*_*α*_) of dietary component α, denoted as *S*_*iα*_, is defined as the ratio between the deviation in *p*_*i*_ and the perturbation amount in *r*_*α*_, i.e., *Siα* ≡ Δ*p*_*i*_/Δ*r*_*α*_. **b**, The ROC curve based on TP rates and FP rates, obtained by setting different susceptibility thresholds for classifications of species-resource interactions in synthetic data generated from MiCRM, where the ground truth of the diet-microbe interactions are encoded in the resource-utilization matrix *U* = (*U*_*iα*_) with *U*_*iα*_≥ 0 represents the rate of species *i* takes up the resource *α*. The AUROC = 0.635, if we compare the susceptibility matrix *S* = (*S*_*iα*_) with the raw resource-utilization matrix *U* = (*U*_*iα*_). By focusing on those strong interactions, e.g., *U*_*iα*_ ≥ 0.005 (or 0.02), we have AUROC = 0.708 (or 0.735), respectively. **c**, Precision–recall (PR) curves for the same task shown in **b**. The corresponding AUPRC values are 0.364 for *U*_*iα*_ ≥ 0.000, 0.323 for *U*_*iα*_ ≥ 0.005, and 0.176 for *U*_*iα*_ ≥ 0.020. **d**, The susceptibility values for the top 20 microbial species (annotated at the species level with the largest absolute susceptibility values) to the top 10 dietary components exhibiting the strongest overall effects in the DMAS cohort data.

We first validated this idea using the synthetic data generated from MiCRM by examining whether the inferred susceptibility matrix *S* captures the underlying resource-utilization matrix *U* encoded in the MiCRM. We quantified concordance between predicted susceptibilities and ground-truth interactions using the AUROC (Area Under the Receiver-Operating Characteristic curve). Across all interactions, we achieved AUROC = 0.635, largely reflecting the intrinsic difficulty or recovering numerous weak interactions (see **Fig.6b**). Restricting the analysis to strong interactions markedly improved performance (AUROC=0.708 for *U*_*iα*_ ≥ 0.005; AUROC=0.735 or *U*_*iα*_ ≥ 0.02). These results indicate that a well-trained MLP can indeed infer species-resource consumption interactions, especially for those strong interactions. Given the strong sparsity of the interaction matrix, we further evaluated performance using the Area Under the Precision-Recall Curve (AUPRC), which is more sensitive to class imbalance. The model achieved AUPRC = 0.364 across all interactions (i.e., *U*_*iα*_ > 0), with AUPRC = 0.323 and 0.176 at thresholds *U*_*iα*_ > 0.005 and > 0.02, respectively. Notably, these values substantially exceed the low baseline AUPRC values (i.e., the positive class prevalence): 0.203, 0.105, and 0.027 for *U*_*iα*_ threshold of 0, 0.005, and 0.02, respectively, indicating enrichment of true interactions among top-ranked predictions. The decrease in AUPRC at higher thresholds reflects the increasing rarity of strong interactions. Together, these results indicate that the susceptibility analysis of a well-trained MLP can recover species-resource consumption interactions, particularly prioritizing stronger interactions despite the highly imbalanced setting.

Then, we applied the susceptibility analysis to the real data in the DMAS cohort (**Fig.6c**). This analysis revealed several coherent patterns. For example, we found positive susceptibility of multiple *Bacteroides* species (including *Bacteroides stercoris, B. dorei, B. vulgatus*, and *B. ovatus*) following perturbations induced by readily bioavailable nutrient sources, such as nutritional drinks and powdered nutritional supplements. This is consistent with the well-established ecological strategy of *Bacteroides* as metabolically flexible generalists, which are capable of rapidly utilizing diverse and readily bioavailable substrates, including simple carbohydrates, processed polysaccharides, and host-accessible nutrients that are typically enriched in processed foods^35–37^. Moreover, we found that key butyrate-producing taxa, e.g., *Faecalibacterium prausnitzii, Alistipes putredinis*, and *Roseburia* showed positive susceptibilities to plant-based foods such as nuts, seeds, and guacamole, as well as to protein-rich dairy products, while exhibiting weak or negative susceptibilities to highly processed foods. This pattern reflects the specialized metabolic requirements of these obligate anaerobic fermenters, which depend on complex carbohydrates, resistant starches, and cross-feeding metabolites generated during fiber fermentation^38,39^. Together, the susceptibility analysis of real data indicates that a well-trained MLP can indeed capture biologically meaningful interactions between specific dietary components and microbial taxa.

In addition to the susceptibility analysis, we also applied SHapley Additive exPlanations (SHAP^40^) analysis to explain the prediction of MPDL on the diet recommendations for single-species enhancement. We found that resources 44 and 28 had the highest mean absolute SHAP values across all samples, indicating that they were the most important features in MPDL (see **Fig.S8a**). Interestingly, those two resources are preferably consumed by target species in MiCRM. We also compared the difference between pre-intervention and post-intervention diet, finding that the resources preferably consumed by target species, e.g., resources 44, 24, 28, and 26, were substantially increased, while those resources that were rarely consumed by target species were decreased (see **Supplementary Table 1**). This strong alignment between SHAP-inferred dietary drivers and the ground-truth species-diet interactions indicates that MPDR effectively learns the underlying microbe-diet relationships and is also robust to the different connectivity (**see Fig.S8b, Supplementary Table 1**).

## Discussion

The human intestine harbors trillions of microbial species. Dysbiosis of the gut microbiome has been associated with many diseases. Diet is considered to be a safe ‘engineer’ of the human gut microbiome^41^. In this work, we proposed a machine learning-based method to prescribe personalized diet recommendations to steer the microbiome toward desired states. Our conceptual framework aligns with recent human dietary-microbiome intervention studies such as the microbiota-directed complementary food (MDCF-2)^42^ trials and metabolite-based dietary signature work (e.g., Shinn et al.)^43^. These studies demonstrated that engineered dietary formulations or food-linked microbial/metabolite signatures can reproducibly modulate or predict gut microbial states in real-world settings. Their findings reinforce the biological plausibility of using diet as a controllable lever to shape microbiome configurations, thereby supporting the central premise underlying MPDR.

We admit that the current work has several limitations. First, we analyzed the DMAS dataset effectively as cross-sectional data. Directly incorporating longitudinal dietary trajectories rather than treating samples as independent may further improve the model’s ability to capture temporal diet-microbiome interactions and enhance prediction accuracy. Second, we validated MPDR using the DMAS data by comparing the MLP-predicted post-intervention compositions and the target compositions. A more rigorous validation of MPDR will require prospective diet intervention studies where the post-intervention compositions are measured, instead of predicted by any machine learning models. Third, this study is limited by the lack of external validation across independent cohorts. Currently, very few publicly available datasets include paired baseline microbiome, detailed dietary intake, and post-intervention microbiome measurements. Future work should evaluate model generalizability across diverse populations and dietary contexts as such datasets become available. Additionally, we didn’t leverage phylogenetic relationships of microbial species in our machine learning models. Incorporating phylogenetic relationships, e.g., through the PhILR transformation^44^, may further enhance the performance of composition prediction and/or diet recommendation. Another limitation is that models trained on observational diet-microbiome datasets may preferentially capture associations involving well-characterized and prevalent taxa, potentially reflecting biases in measurement, annotation, and prior literature. This raises the possibility that part of the predictive signal recapitulates known associations rather than fully generalizable interaction rules. More rigorous benchmarking frameworks incorporating curated positive and negative interaction sets^45^, or biologically informed synthetic data generation (e.g., metabolic model-constrained simulations^46^), may help address this limitation in future work.

A key assumption underlying our framework is that a given species assemblage-diet configuration maps to a unique microbial composition. While this assumption holds in fully observed systems such as MiCRM simulations, where the complete resource environment is specified, it might not hold for real-world data. Dietary representations derived from databases, e.g., Food and Nutrient Database for Dietary Studies, capture only a limited subset of biologically relevant exposures. Various factors, including food matrix effects, industrial additives, cooking methods, and environmental contaminants, introduce latent variables that are not reflected in the observed dietary profile. Consequently, identical species assemblage-diet pairs may correspond to multiple microbial compositions. In this context, our framework should be interpreted as learning an effective mapping that approximates the expected microbiome state conditioned on the observed inputs, rather than a strictly deterministic one-to-one relationship. Importantly, this limitation arises from incomplete measurement of dietary exposures rather than from the modeling framework itself. Future work may improve identifiability by incorporating richer dietary representations, including food processing metadata, additive exposure profiles, or metabolomics-derived dietary proxies. In addition, probabilistic extensions that explicitly model latent dietary variables could provide a principled way to quantify uncertainty in inferred microbial compositions and further refine predictive performance.

## METHODS

### Synthetic data generation

To validate our framework, we leverage an existing Microbial Consumer Resource Model (MiCRM)^26^. The dynamics of species and resources are given by the following ordinary differential equations:

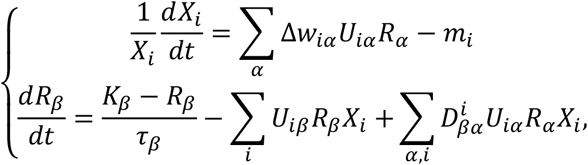

where *X*_*i*_ is the abundance of species *i* and *R*_*α*_ is the abundance of resource *α*. The entry *U*_*iα*_ represents the rate of species *i* takes up the resource *α*. 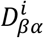 denotes the number of molecules of resource *β* secreted by the environment species *i* per molecule of resource *α* it uptakes. We introduced a parameter *C* to control the connectivity of matrix *U*, e.g., *U*_*iα*_ = 0 with probability 1 − *C*. Δ*w*_*iα*_ is the net biomass yield (growth gain) per unit of resource *α* consumed by species *i*, and *m*_*i*_ is the species-specific maintenance/mortality rate.

For the resource dynamics, *K*_*β*_ is the external supply (set-point) of resource *β* and *τ*_*β*_ is its turnover (replenishment) time constant. The consumption term ∑_*i*_ *U*_*iβ*_ *R*_*β*_ *X*_*i*_ aggregates removal of resource *β* by all consumers. The cross-feeding term 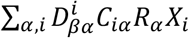 accounts for metabolic byproducts: 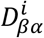 is the stoichiometric coefficient giving the number of molecules of resource *β* secreted by species *i* per molecule of resource *α* it processes, and *C*_*iα*_ (dimensionless) modulates the effective consumption/processing propensity of species *i* on resource *α*. Indices run over species *i*=1,…,*N* and resources *α, β*=1,…,*M*.

The species collection of each sample is a selected sample of 80 consumers from the meta species pool of 100: 50 species were shared across all samples, and the remaining 30 were randomly selected from the other 50 species in the pool. The scale approximates within-host richness in the human gut microbiome (about 100-200 bacterial species per individual)^47^. To avoid bias toward a specific ecological configuration while preserving realistic metabolic structure, the resource-utilization matrix *U* was explicitly specified within the MiCRM framework based on four metabolic families, reflecting the well-established organization of microbiome communities into a limited number of metabolic guilds. Each metabolic family was assigned a characteristic resource-uptake prior in MiCRM, from which species-level utilization profiles were generated and subsequently sparsified using a Bernoulli connectivity mask to control network density. This procedure produces structured, non-random resource uptake patterns arising from model design rather than arbitrary random sampling. The species secretion matrix *D* was likewise defined within the MiCRM to yield stoichiometrically consistent cross-feeding, thereby introducing realistic metabolic byproduct interactions among species. We set the total number of resources to *M* = 60, matching the dimensionality of standard dietary surveillance systems that typically track ~60-70 nutrient variables^48^.

To evaluate universality and robustness, we systematically varied a set of structural/experimental-design hyperparameters of the MiCRM. Specifically, we swept the perturbation amplitude *σ*∈{0.02, 0.05, 0.1}, consumer specialism level *g*∈{0.01, 0.1, 0.5}, network connectivity *C*∈{0.2, 0.4, 0.6}, and train/test sampling ratio r∈{0.5, 1, 2, 5, 10}. For each (*σ, g, C*, r) combination, we performed 10 independent folds (fold=1,…,10) with fresh random draws of species sets and resource profiles. For a given fold, we generate a large pool of samples and split them into train and test by n_train = round (n_pool · r / (1 + r)), leaving n_test = n_pool − n_train.

To systematically examine the performance of MPDR in terms of universality, we first generated a baseline external resource supply *q*_*b*_ drawn from a uniform distribution on (0,1), then normalized so that the total resource amount is 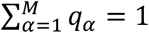. This will remove the arbitrary scale so the model learns relative resource distribution rather than simulation-imposed magnitude. In our implementation, environmental perturbations were introduced through the MiCRM parameter constructor (mcrm_params), which generates resource supply configurations whose variability is controlled by the perturbation amplitude *σ*. This procedure generates resource profiles that differ from one another in a controlled manner, with the magnitude of variation determined by the perturbation amplitude. We ran the MiCRM dynamics to the steady state to obtain the baseline community with species assemblage ***z***. Then, a second independent perturbation of the same form was subsequently applied on top of the already perturbed resource profile, and the system was re-integrated starting from ***z***, yielding the resulting community ***p***.

### MPDR framework

The key idea of our MPDR framework is to use a machine learning model to predict microbial compositions from species assemblages and dietary intake profiles. Given a species pool Ω = {1, ⋯, *N*} associated with a particular habitat (e.g., the human gut), any microbiome sample collected from this habitat can be considered as a local community assembled from the species pool Ω. Consider a set of local communities collected from this habitat, e.g., gut microbiome samples from different hosts. In practice, the species assemblage of a sample can be represented by a binary vector ***z*** ∈ {0,1}^*N*^, where its *i*-th entry satisfies ***z***_***i***_ = 1 (or 0) if species *i* is present (or absent) in this sample. This is motivated by considerations of robustness, because relative abundance measurements in microbiome data are inherently noisy, particularly for low-abundance taxa, and are further subject to compositional constraints. To demonstrate this point, we evaluated the robustness of our modeling framework by introducing controlled measurement noise into baseline abundance profiles and systematically compared predictive performance using binary inputs (i.e., species presence/absence patterns)_;_ versus continuous relative abundance inputs. Under increasing noise levels, binary inputs consistently yielded more stable and improved predictive accuracy, supporting its use as a robust representation of the baseline microbiome state (see **Fig.S9)**. The dietary intake profile of a sample can be represented by a vector 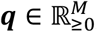, where its *i*-th entry is the abundance of food item *i* in the dietary intake data. The microbial composition of this sample supplied by ***q*** is characterized by a vector ***p*** ∈ Δ^*N*^, where its *i*-th entry ***p***_***i***_ is the relative abundance of species *i* in this sample. Here, instead of using any population dynamics model in community ecology, we used Multi-Layer Perceptron^49^ (MLP) to learn the map 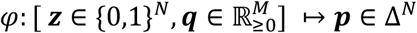 directly from a set of microbiome samples from a habitat. Notley, both *N* (the number of species) and *M* (the number of dietary features) are automatically determined by the dataset rather than manually fixed. Specifically, for each dataset, we determine *N* by the total number of detected species across all samples, and *M* by the number of dietarty features provided. This ensures that the model output layer always corresponds to the entire observed species pool, avoiding the use of any restricted sub-composition.

To learn such a mapping, we concatenate the dietary intake profile ***q*** with the baseline microbiome vector ***z***, forming the joint input [***q, z***] ∈ ℝ^***M***+***N***^. This vector is then passed into a shared two-layer fully connected network to generate the Dirichlet concentration parameters of the target microbial composition:

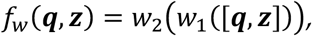

where *w*_1_ ∈ ℝ^256⨯(*M*+*N*)^, *w*_0_ ∈ ℝ^*N* ⨯256^ to learn more complex patterns in the data. A softplus transformation is applied to ensure positivity of the Dirichlet parameters, with a small constant (10^−6^) added to enforce a strict lower bound and improve numerical stability during training:

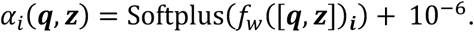

We further apply an element-wise species-presence mask *m*_*i*_ = 1(***z***_*i*_ > 0), ensuring that species absent in the baseline assemblage do not receive non-negligible concentration parameters. Specifically, for absent species (*m*_*i*_ = 0), the corresponding Dirichlet parameters are set to a small positive constant (10^−6^) to maintain a valid Dirichlet distribution:

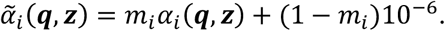

The predicted composition is then obtained by normalizing the masked concentration parameters:

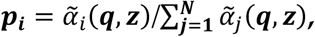

which ensures that the total abundance sums to one and that species absent in the assemblage (i.e., *z*_*i*_ = 0) receive zero predicted abundance.

The MLP is trained to minimize the following loss:

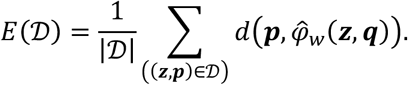

Here 𝒟 is the dataset, and *d* is the masked Dirichlet negative log-likelihood loss. We use a Dirichlet distribution because microbial community compositions are compositional data constrained to the simplex, and the Dirichlet likelihood provides a principled probabilistic model defined directly on this space, capturing relative abundances and inter-taxon dependencies without requiring transformation^50^. Model parameters are learned by minimizing the negative log-likelihood, which corresponds to maximum likelihood estimation of the predicted distribution and is a standard objective for probabilistic neural networks with parametric output distributions^51^. The masked formulation allows the loss to ignore absent or unobserved taxa, improving robustness to sparsity. We used a batch size of 20 and trained for 1,000 epochs for all datasets. Model optimization was performed using the Adam optimizer with a learning rate of 0.01.

To further compare the representational power of different architectures, we designed an additional Transformer-based model that shares the same unified MLP prediction head but differ in their front-end feature extraction modules. This model applies two independent Transformer encoders to the dietary features and the baseline microbiome composition, respectively. Prior to encoding, both species and diet tokens are globally ordered according to their abundance, estimated from the training microbiome composition ***p*** and the dietary feature ***q*** ^28^, thereby introducing an explicit abundance-based inductive bias before feature fusion. The resulting embeddings are then fused to capture joint dependencies between dietary intake and species context. For this Transformer model, the fused front-end representations are concatenated with the baseline microbiome vector and passed into a shared two-layer fully connected MLP prediction head, which outputs the Dirichlet concentration parameters for the target microbial composition. This design ensures that the comparison across architectures focuses solely on their representational power, while keeping the output mapping consistent.

In addition to these neural architectures, we also benchmark three classical regression baselines, multi-output SVR, RF regression, as well as XGBoost regression, which operate directly on the concatenated tabular features [***z*** ∣ ***q***] and produce continuous abundance predictions for each taxon. Following the implementation, these raw outputs are post-processed by clipping negative values and renormalizing each row to sum to one, ensuring valid microbial composition vectors ***p***.

All models were evaluated using the same train/test splits, input/output definitions, and evaluation metrics to ensure fair comparison across architectures. Hyperparameters for traditional machine learning models (SVR, RF, and XGBoost) were independently optimized based on validation set performance.

### MPDR framework - dietary recommendation

After the training of MLP to predict microbial composition from species assemblage and dietary intake, the personalized dietary recommendation was achieved through gradient-based (GB) optimization of the dietary intake profile. Given any dietary intake profile ***q***, baseline species collection ***z***, and the regulation goal, e.g., the dissimilarity between the recommended-diet projected community composition and the target composition, or relative abundance of target species, we defined a loss function *L* to quantify the mismatch. For each sample, the optimization proceeds by updating a set of unconstrained logits. (For synthetic data generated by MiCRM, the diet or resource profiles are compositional so the logits are further mapped to a valid resource profile via a non-negativity and simplex projection.) At each iteration, the logits are updated using gradients of the loss function. This process is repeated for a fixed number of optimization steps (default: 1000), with early stopping when no further improvement is observed.

For each individual, the dietary intake profile ***q*** was initialized from existing dietary records or set to zero, and iteratively optimized to minimize the Bray-Curtis dissimilarity between the model-projected microbial composition and the target state. The optimization objective is defined as:

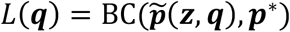

where BC(⋅,⋅) denotes the Bray-Curtis dissimilarity, 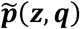 denotes the microbial composition predicted by the trained model for a given (***z, q***) pair, and ***p***\* represents the target microbial state. This procedure yields an optimized diet ***q***\*that minimizes dissimilarity between predicted and target microbiome states.

An alternative method to achieve optimal diet recommendation is based on the simulated annealing (SA) algorithm: (1) randomly perturb the current dietary intake profile ***q***. (2) calculate new loss function denoted as *L*_IJK_; (3) update dietary intake profile as ***q***_new_ if *L*_new_ < *L*_current_ or with probability exp (−(*L*_new_ – *L*_current_)/*T*) if *L*_new_ > *L*_current_. Repeat steps (1)-(3) 5,000 times to obtain the final recommendation ***q***_*r*_, and the temperature was decreased geometrically as *T* =*T*_0_*α*, where *T*_0_ (we choose 100) is the initial temperature and *α* = 0.999.

### DMAS Data Preprocessing

The DMAS dataset provides daily food-intake records and paired gut microbial profiles. All 65 nutrient variables and 1,245 food-intake features were taken directly from the publicly available DMAS dataset^6^. DMAS comprises 34 subjects followed for 17 consecutive days with paired daily dietary records and fecal microbiome samples. Dietary intake was assessed using the automated self-administered 24-hour (ASA24) dietary assessment tool. The cohort included both male and female participants, with expected sex differences in anthropometric measures (e.g., weight, height, and waist circumference) as reported in the original study. The study design also incorporated a 10-day, double-blind dietary intervention comparing medium-chain triglycerides (MCTs) and extra-virgin olive oil (EVOO). Detailed demographic and clinical characteristics of the cohort have been previously described (see the original publication Table 1 and STAR Methods), and are not reproduced here. Dietary entries and microbiome samples were aligned using a unique subject-day identifier, and missing dietary values were set to zero. The microbiome table originally contained 223 species; raw counts <130, corresponding to a relative abundance of approximately 10^⁻6^, were set to zero, and species present in <15% of samples were removed, resulting in 138 retained species and 50 genera (**Supplementary Table 2)**. Consecutive-day observations were used to construct training and testing pairs. A randomized control dataset was additionally generated by shuffling the 500 most frequently consumed foods within each sample.

To construct training and testing pairs, we used longitudinal information within the cohort. For each individual, consecutive-day observations (RecordDayNo + 1) were used to form training pairs of baseline microbial composition ***z***, next-day composition ***p***, and dietary intake ***q***. Testing pairs were produced analogously, including all cases exhibiting measurable changes in microbial profiles. In addition, species/genus-presence vectors were generated by binarizing microbial compositions (0/1), providing the collection vector required by the MPDR framework. The model training and the diet recommendation procedure are kept consistent with those used in the previous simulation data. Specifically, we employ an MLP model that takes the concatenated diet and microbiome features as joint inputs, and diet optimization is performed using the network-gradient-based approach.

## Data availability

The DMAS dataset is available from https://github.com/knights-lab/dietstudy_analyses.

## Code availability

Python code used in this work is available at https://github.com/danhuangagu-ship-it/MPDR.

## Acknowledgements

Research reported in this publication was supported by grants R01AI141529, R01HD093761, RF1AG067744, UH3OD023268, U19AI095219, and U01HL089856 from National Institutes of Health. X.-W.W. acknowledges the funding support from National Institutes of Health (K25HL166208).

## Author Contributions

Y.-Y.L. conceived and designed the project. X.-W.W. and D.H. performed all the numerical calculations and real data analyses. X.-W.W., D.H. and Y.-Y.L. analyzed and interpreted the results. X.-W.W., D.H. and Y.-Y.L. wrote the manuscript. All authors edited the manuscript.

