## Supplemental figures for "Machine learning-based Personalized Dietary Recommendations to Achieve Desired Gut Microbial Compositions"

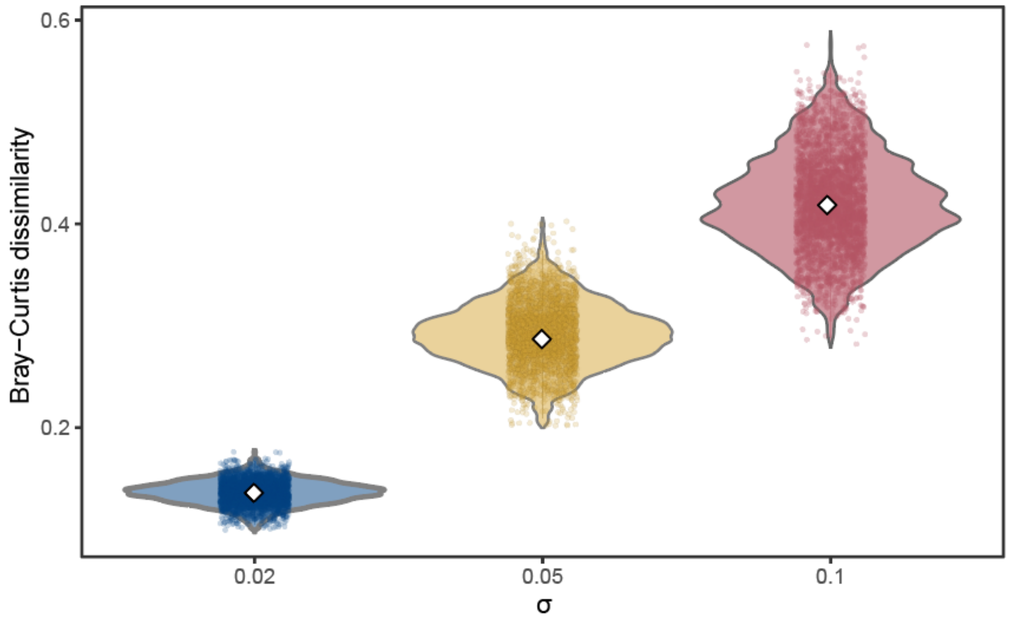


**Figure S1**: **Bray-Curtis dissimilarity between each simulated diet and the mean diet within its corresponding σ group ($\sigma = 0.02$, 0.05, 0.1).** Each box shows the distribution of dietary variability generated under different perturbation levels, illustrating how σ controls the magnitude of diet fluctuations used in the simulation framework.

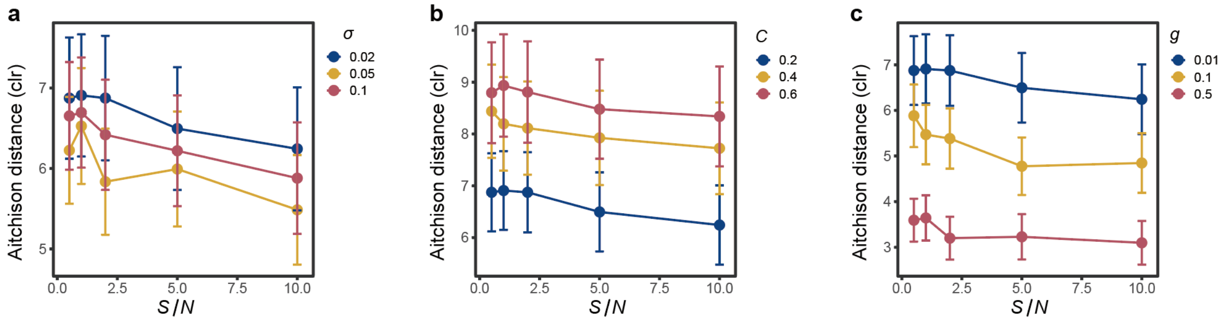


**Figure S2: Prediction error quantified as Aitchison distance between the true and predicted compositions**. The error bar represents the mean of standard deviation over 10 random splitting datasets. The x-axis encodes the training sample ratio ($\frac{S}{N}$). In panel a, $C = 0.2$ and $g = 0.01$. In panel b, $\sigma = 0.02$ and $g = 0.01$. In panel c, $\sigma = 0.02$ and $C = 0.2$.

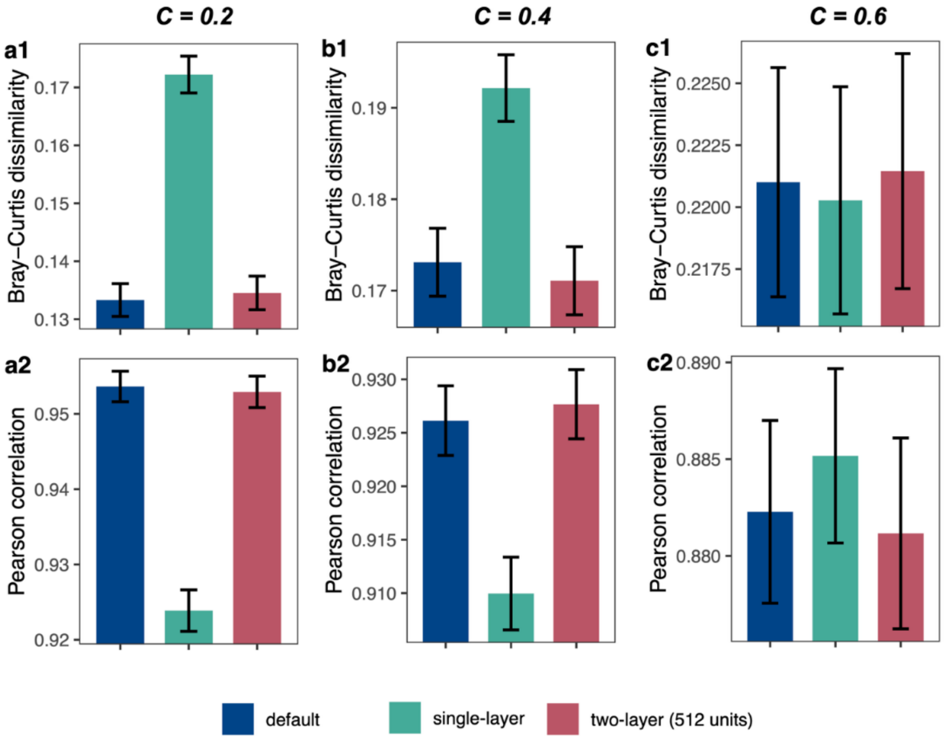


**Figure S3: Performance comparison of various MLP architectures across different connectivity levels.** Columns correspond to different connectivity settings, and rows report performance metrics (top: Bray–Curtis dissimilarity; bottom: Pearson correlation). Results are shown for different MLP architectures, including a two-layer MLP with hidden units 256 (default), a single-layer MLP with hidden units 256 (single-layer), and a two-layer MLP with hidden units 512.

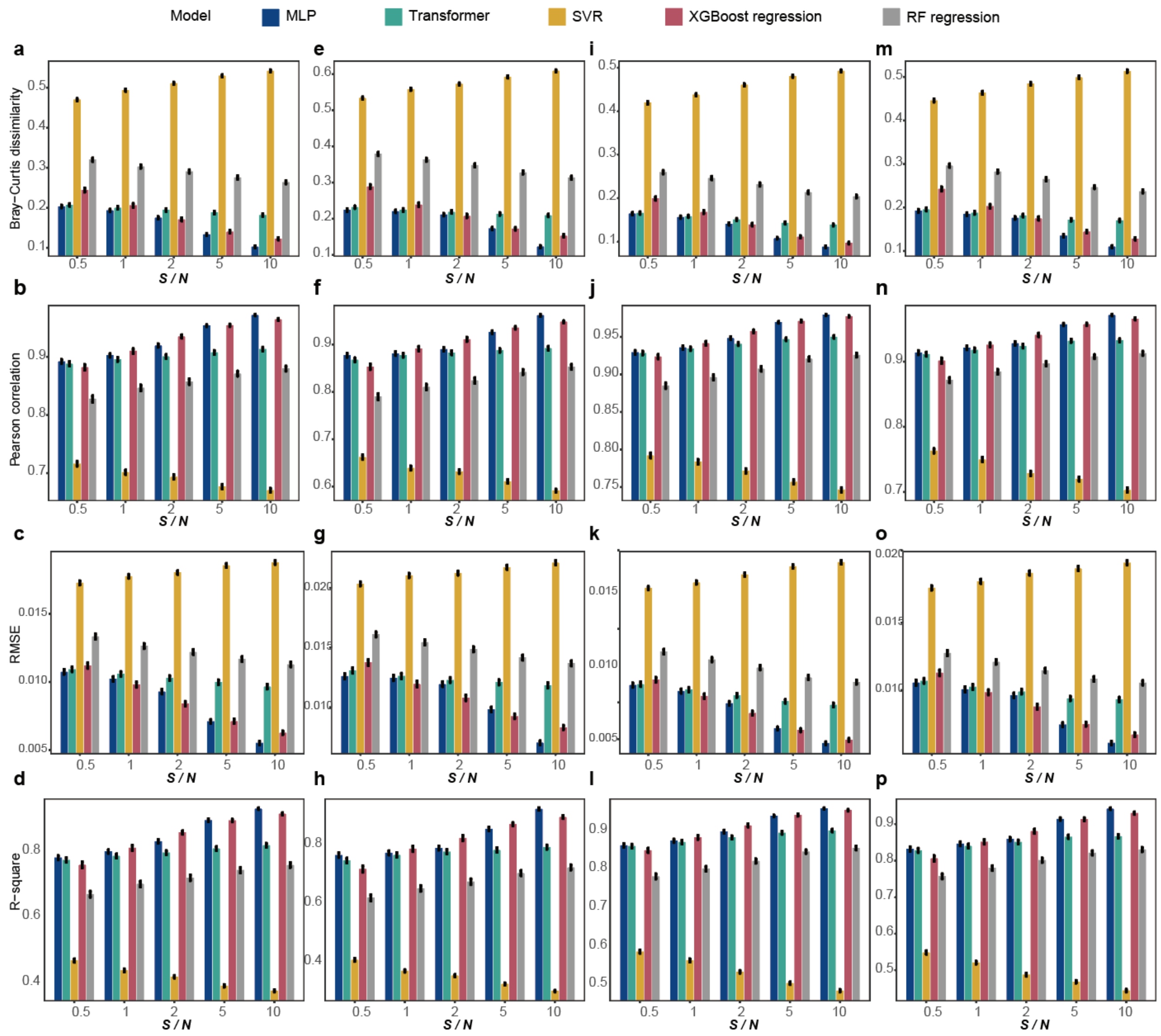


**Figure S4: Benchmarking the performance of five machine learning models (MLP, Transformer, SVR, XGBoost regression, and RF regression) in microbial composition prediction.** Prediction performance is quantified by (a,e,i,m) Bray-Curtis dissimilarity between the ground-truth compositions computed by the MiCRM and the compositions predicted by the five machine learning models (the lower the better); (b,f,j,n) Pearson correlation between the ground-truth species abundances and the predicted ones (the higher the better); (c,g,k,o) RMSE (the lower the better); and (d,h,l,p) R-square (the higher the better). In the MiCRM simulations, we have a species pool of N=100 and a resource pool of M = 60. For each parameter combination, S samples were generated to train the models, and ten random train/test splits were used for evaluation. Different panels correspond to different parameter settings in MiCRM: (a,b,c,d) $\sigma = 0.1$, $C = 0.2$ and $g = 0.01$, (e,f,g,h) $\sigma = 0.1$, $C = 0.4$ and $g = 0.01$, (i,g,k,l) $\sigma = 0.05$, $C = 0.2$ and $g = 0.01$, (m,n,o,p) $\sigma = 0.05$, $C = 0.4$ and $g = 0.01$. The x-axis represents the relative training sample size ($\frac{S}{N}$).

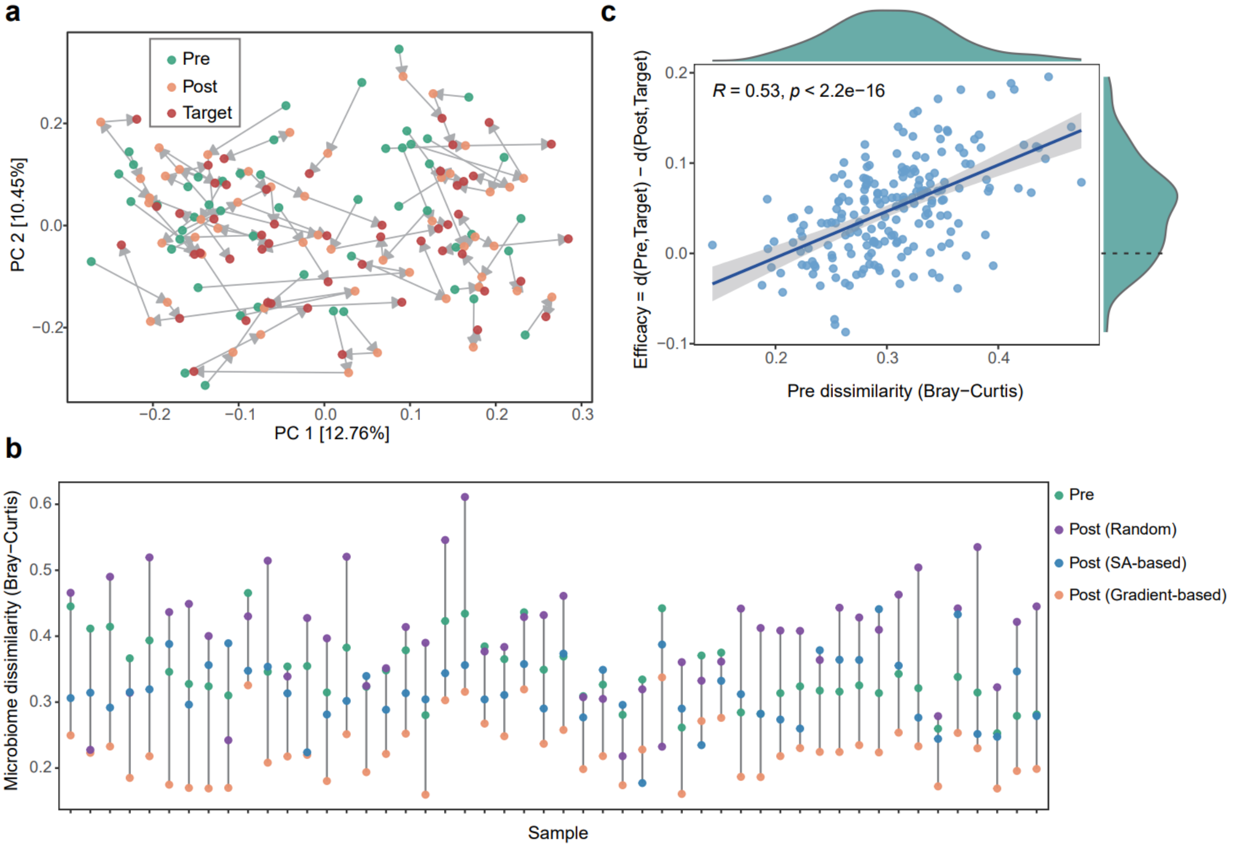


**Figure S5: In silico validation of dietary recommendation in MPDR framework.** Results are obtained for the pools of $N = 100$ species and 60 resources with a microbiome consumer-resource model. We generated 500 samples to train MPDR and an additional 200 samples to validate the recommendation of MPDR. For each validation sample with target microbial composition $\boldsymbol{p}^*$, we randomly perturbed its nutrient profile $\boldsymbol{q}_{\mathrm{H}}$ into another nutrient profile $\boldsymbol{q}_{\mathrm{U}}$ and associated baseline microbial composition $\boldsymbol{p}_{\mathrm{U}}$. MPDR was applied to find the optimal nutrient profile $\boldsymbol{q}^*$ that minimizes dissimilarity between recommended-diet projected composition $\tilde{\boldsymbol{p}} = \varphi(\boldsymbol{z}, \boldsymbol{q}^*)$ and $\boldsymbol{p}^*$. Then, ran MiCRM again by using $\boldsymbol{q}^*$ as the nutrient profile to obtain the new or recommended-diet projected composition $\tilde{\boldsymbol{p}}$. **a**, PCoA plot showing the microbial composition dissimilarity between baseline compositions ($\boldsymbol{p}_U$), recommended-diet projected ($\tilde{\boldsymbol{p}}$) and target compositions ($\boldsymbol{p}^*$). **b,** Microbiome dissimilarity (Bray-Curtis) between the baseline microbiome composition ($\boldsymbol{p}_{\mathrm{U}}$) and the target composition ($\boldsymbol{p}^*$) (Pre, green), and between the post-intervention microbiome compositions ($\tilde{\boldsymbol{p}}$) obtained using different diet recommendation strategies: random, simulated annealing (SA), and gradient-based optimization, and the target composition ($\boldsymbol{p}^*$)(Post). In all panels, $\sigma = 0.1$, $C = 0.4$ and $g = 0.01$. **c,** Recommendation efficacy $d(\boldsymbol{p}_{\mathrm{U}}, \boldsymbol{p}^*) - d(\tilde{\boldsymbol{p}}, \boldsymbol{p}^*)$ versus the pre-dissimilarity $d(\boldsymbol{p}_{\mathrm{U}}, \boldsymbol{p}^*)$. In panels a and b, we only showed 50 samples with the highest recommendation efficacy.

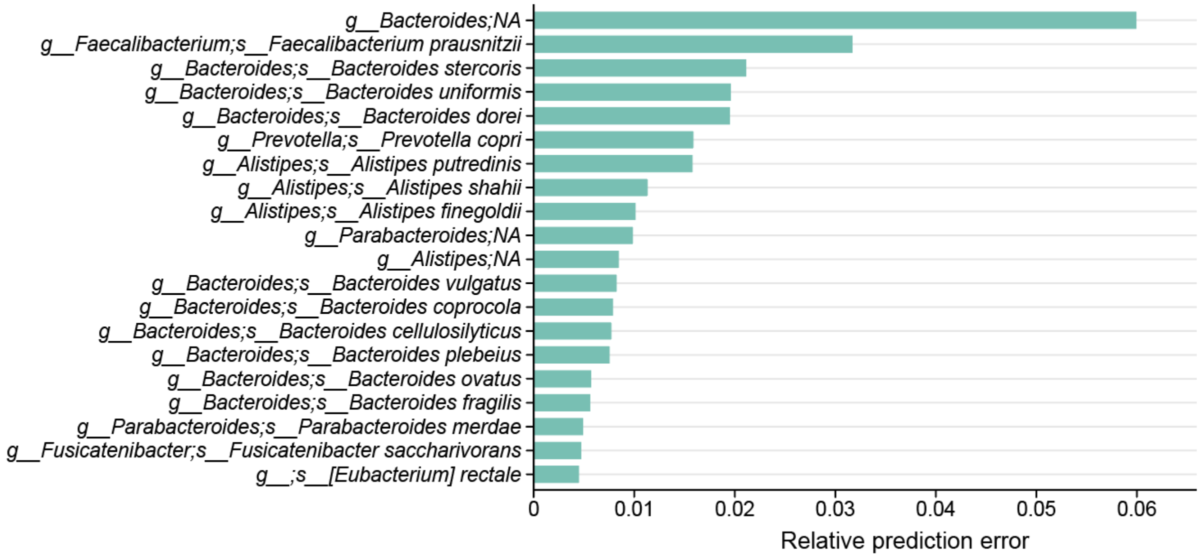


**Figure S6: The 20 taxa with the lowest prediction error in the MPDR model.** Shown are the top 20 microbial taxa ranked by their relative prediction error, defined as the mean absolute prediction error normalized by the mean true relative abundance across all samples. Taxa with very low baseline abundance (<0.001) were excluded prior to ranking to ensure stable and interpretable error estimates. Lower values indicate taxa whose compositions are more accurately captured by the model and are therefore more stably predictable under dietary perturbations.

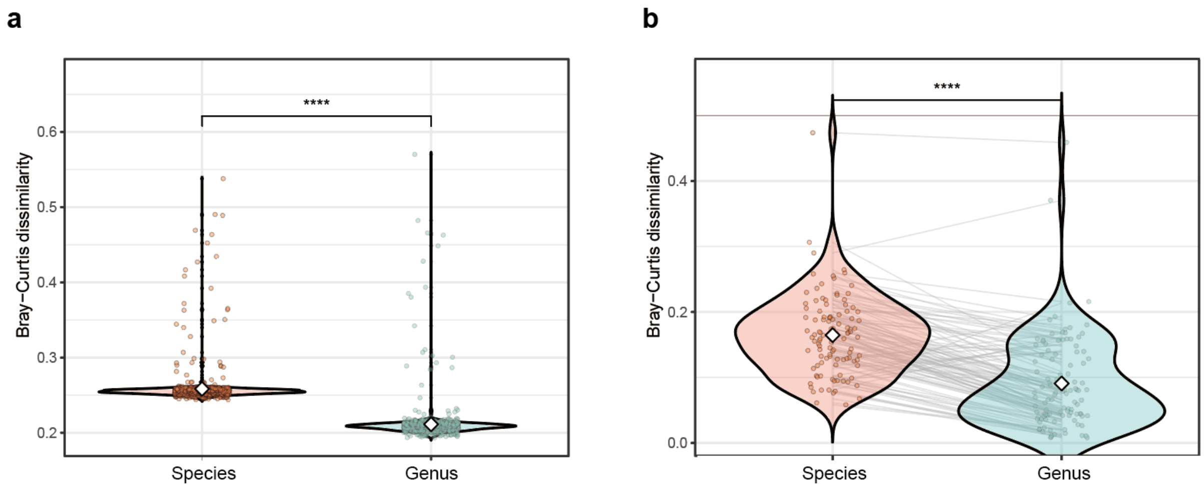


**Figure S7: Comparison of the MPDR framework for community-wise regulation at the species and genus levels using the DMAS dataset.** Prediction errors were quantified using Bray-Curtis dissimilarity between predicted and ground-truth microbiome compositions during the validation phase (**a**; n = 1000 epochs), and Bray-Curtis dissimilarity between the desired microbial composition and Model-predicted endpoints under recommended diets (**b**; n = 107 samples), respectively. For each test sample, prediction errors from the two models (Species-aware and Genus-aware) were compared pairwise, and statistical significance was assessed using paired Wilcoxon signed-rank tests. ****: p-value < 0.00001.

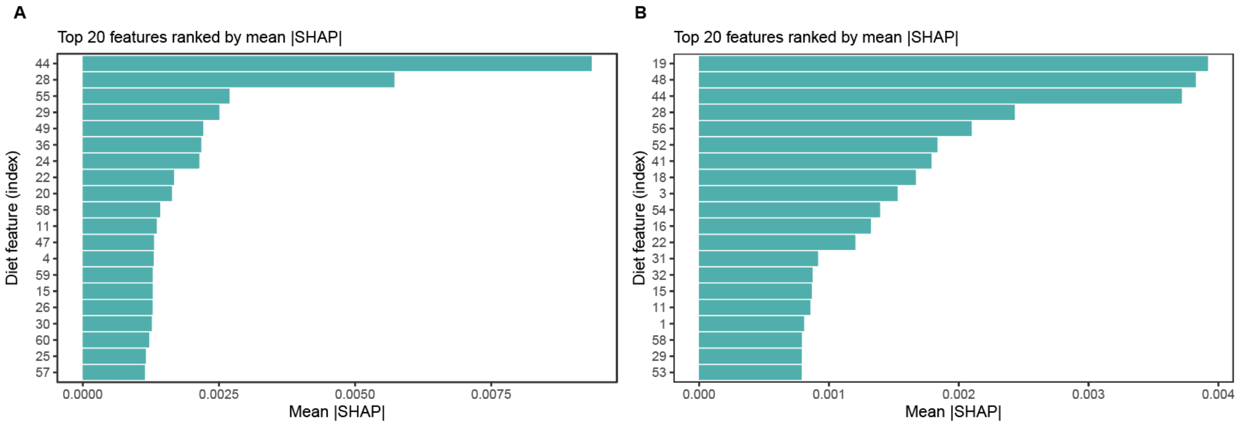


**Figure S8: Interpretation of in silico validation of dietary recommendation in MPDR framework for species-wised regulation.** Results are obtained for the pools of $N = 100$ species and 60 resources with a microbiome consumer-resource model. We generated 500 samples to train MPDR and an additional 200 samples to validate the recommendation of MPDR. For each validation sample, we randomly perturbed its nutrient profile $\boldsymbol{q}_{\mathrm{H}}$ into another nutrient profile $\boldsymbol{q}_{\mathrm{U}}$ and associated baseline microbial composition $\boldsymbol{p}_{\mathrm{U}}$. MPDR was applied to find the optimal nutrient profile $\boldsymbol{q}^{*}$ that minimizes relative abundance of target species $x_t$.Then, ran MiCRM again by using $\boldsymbol{q}^{*}$ as the nutrient profile to obtain the new or recommended-diet projected composition $\tilde{\boldsymbol{p}}$. **a-b,** SHAP analysis of MPDR dietary recommendations. Panel **a** shows the top 20 dietary features (ranked by mean absolute SHAP value) contributing to the abundance change of species 22 under $C = 0.2$. Panel **b** shows the corresponding top 20 dietary features influencing species 22 under $C = 0.4$. In two panels, $\sigma = 0.1$, and $g = 0.01$.

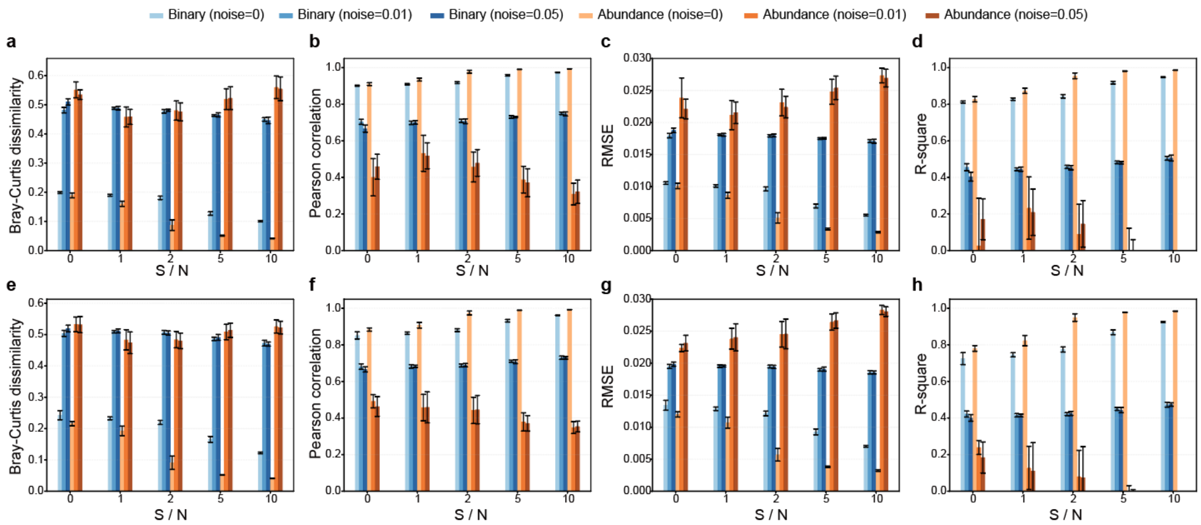


**Figure S9. Benchmarking the performance of MLP models using two different input types: binary (i.e., species presence/absence pattern) vs. continuous (species relative abundances) under increasing additive Gaussian noise levels for microbial composition prediction.** Prediction performance is quantified by (a,e) Bray-Curtis dissimilarity between the ground-truth compositions computed by the MiCRM and the compositions predicted by machine learning model (the lower the better); (b,f) Pearson correlation between the ground-truth species abundances and the predicted ones (the higher the better); (c,g) RMSE (the lower the better); and (d,h) R-square (the higher the better). The binary-input model was trained on presence/absence data derived from abundance measurements after noise addition (threshold >0 for presence, ≤0 for absence), while the abundance-input model was trained on normalized compositional abundance data after noise addition. Additive Gaussian noise with standard deviation σ_noise = {0, 0.01, 0.05} was applied to the original abundance data before thresholding (binary mode) or renormalization (abundance mode). In the MiCRM simulations, we have a species pool of N=100 and a resource pool of M = 60. For each parameter combination, S samples were generated to train the models, and ten random train/test splits were used for evaluation. Different panels correspond to different parameter settings in MiCRM: (a-d) $\sigma = 0.1,\ C = 0.2$ and $g = 0.01$, (e-h) $\sigma = 0.1,\ C = 0.4$ and $g = 0.01$. The x-axis represents the relative training sample size ($\frac{S}{N}$).
